# Preclinical efficacy and safety of Tegavivint in Wnt-activated hepatocellular carcinoma

**DOI:** 10.64898/2026.07.14.738585

**Authors:** Toshiyasu Suzuki, Ciara Curran, Tom Drake, Stephanie May, Kyi Lai Yin Swe, Anastasia Georgakopoulou, Megan Quince, Fiona Chalmers, Emma Paterson, Aundrietta Duncan, Stephen Horrigan, Mark Eamonn Kelly, Colin Nixon, Victor Hugo Villar, Thomas Graham Bird

**Affiliations:** Cancer Research UK Scotland Institute, Glasgow, G61 1BD, UK; School of Cancer Sciences, University of Glasgow, Glasgow, G61 1QH, UK; Centre for Medical Informatics, University of Edinburgh, Edinburgh, EH16 4UX; Iterion Therapeutics, INC., Houston, TX, United States of America; School of Medicine, University of St Andrews, North Haugh, KY16 9TF, UK; Institute for Regeneration and Repair, University of Edinburgh, Edinburgh, EH16 4UU, UK; Cancer Research UK Scotland Centre, Glasgow & Edinburgh, UK

**Author notes:** Centre for Cancer Immunotherapy and Immunobiology, Graduate School of Medicine, Kyoto University, Kyoto, Japan (Toshiyasu Suzuki).

## Abstract

**Background & Aims:** Hepatocellular carcinoma (HCC), a predominant form of liver cancer, remains a significant clinical unmet need. Given that 30–50% of HCC cases harbour mutations in the Wnt/β-catenin signalling pathway, targeting this cascade represents a promising therapeutic strategy. However, the clinical translation of Wnt inhibitors has been hindered by severe adverse events observed in preclinical models and early-phase clinical trials, primarily due to the essential role of Wnt signalling in maintaining normal tissues such as the intestine and bone.

**Methods:** We examined the efficacy of Tegavivint, a first-in-class Wnt pathway inhibitor that targets TBL1, against HCC to elucidate its underlying mechanism of action. We evaluated the dose-response of Tegavivint and its effects on the cell cycle, apoptosis, and Wnt target gene expression using HepG2, HUH6, and HUH7 cell lines *in vitro*. Furthermore, we employed an orthotopic xenograft transplant model using HepG2 cells in immunodeficient mice to assess the safety profile and on-target anti-cancer efficacy of Tegavivint *in vivo*.

**Results:** Tegavivint exhibited potent Wnt pathway-suppressing effects in cancer cells with constitutive Wnt pathway activation. Notably, Tegavivint displayed robust anti-tumour activity across a broad range of HCC cell lines, regardless of their Wnt pathway activation status. While Tegavivint inhibited the Wnt pathway and triggered the activation of apoptotic pathways in most cell lines, our findings suggest that it also can induce cell death by activating alternative non-apoptotic pathways in apoptosis-resistant cancer cells. In an orthotopic transplant mouse model, Tegavivint significantly downregulated Wnt pathway target genes, inhibited cell proliferation, induced apoptosis and suppressed growth in tumours.

**Conclusions:** Taken together, our data establish a robust foundation for evaluating Tegavivint as a novel therapeutic option, specifically tailored for HCC patients harbouring Wnt-driven hepatic malignancies.

## Introduction

Liver cancer, of which hepatocellular carcinoma (HCC) is the most common type, represents a major area of unmet clinical need^1^. Treatment in the tumour’s early stages can be curative but currently systemic chemotherapy, despite entering an era of combination immunotherapies, still offers a low chance of cure^2^. HCC is a heterogeneous tumour driven by a number of distinct oncogenic pathways^3–5^. Rare forms of the tumour have shown small scale opportunities for treatments that target the driver mutations in HCC^6^. However, the bulk of the driver mutations in HCC have proved difficult to target to date^5, 7, 8^.

The Wnt/β-catenin pathway is an intercellular signalling pathway facilitated by signalling using both Wnt and R-Spondin ligands which together act to stabilise cytoplasmic β-catenin permitting its nuclear translocation and activation of downstream targets^9^. These targets (including LGR5, GS-glutamine synthetase, NOTUM, AXIN2 and Cyclin D) have pleiotropic effects but include the stimulation of proliferation and a distinct metabolic programme in hepatocytes, important for hepatic zonation^10^. While the portal venous zone, where oxygen- and nutrient-rich blood enters, serves as a hub for energy synthesis, the opposite pericentral zone, where processed blood exits, is responsible for detoxification, glycolysis (energy consumption) and bile acid synthesis^11^ and is directly specified by active β-catenin signalling pathway within hepatocytes^12, 13^. In liver cancer, the Wnt/β-catenin pathway is one of the most frequently mutated cancer pathways^4^, with mutations typically occurring in the *CTNNB1* gene that encodes β-catenin. This results in a degradation-resistant protein which then persistently activates the oncogenic signalling^14^. The activation of β-catenin in HCC is associated with steatotic liver disease^15^ and distinct molecular characteristics of the tumour^16, 17^. These include activation of β-catenin targets and also the suppression of immune chemoattractants (chemokines) which may act to suppress immune infiltration and response to immunotherapy^18,19^. Intriguingly, β-catenin mutations are also seen as high-risk features of hepatic adenomas because of their associated high rates of malignant transformation^20^. Recent preclinical work suggests that the strength of β-catenin activation needs to be modulated (e.g. by MAPK/mTOR/IGFBP2 mediated pathway suppression) to facilitate malignant expansion^21^. This may explain why HCC has a preponderance for *CTNNB1* mutations compared to other tumours (e.g. *APC* mutations in colorectal cancer). To date, a series of inhibitors targeting the Wnt/β-catenin pathway, including PORCN inhibitors (i.e. LGK974, ETC-159), CBP/β-catenin inhibitors (i.e. PRI-724, ICG-001), Tankyrase inhibitors (i.e. XAV939, JW55), and TCF/β-catenin inhibitor (CGP049090, PKF-115584), have been developed. However, they have not been shown to be clinically effective in HCC^22–24^. Other mechanisms to target β-catenin mutant HCC have been described (including lipid nanoparticle delivered siRNA) and have shown potential in preclinical models of the disease^25^.

Tegavivint is a novel Wnt/β-catenin inhibitor that inhibits the interaction between β-catenin and Transducin-beta-like protein 1X (TBL1 or TBL1X), and its homolog Transducin-beta-like protein 1-related protein 1 (TBL1R or TBL1XR1). Without stimulation by Wnt ligands, the transcription factor TCF and the co-repressor TLE1 form a repressor complex on the promoter regions of Wnt target genes in the nucleus. Upon ligand-receptor activation, β-catenin is imported into the nucleus. The translocation of β-catenin into the nucleus leads to the formation of a complex with TBL1 or TBL1R, which facilitates the displacement of TLE1 and activates the promoter regions of Wnt target genes^26, 27^. Therefore, TBL1 and TBL1R play an important role in regulating the transcriptional activity of Wnt/β-catenin signalling. TBL1 and TBL1R are frequently overexpressed in various cancers, including solid tumours and hematologic malignancies, where their elevated levels are associated with poor prognosis and advanced disease stages^28^. Preclinical mouse models demonstrated efficacy of Tegavivint *in vivo* in various cancer types, including acute myeloid leukaemia (AML)^29, 30^, multiple myeloma^31, 32^, osteosarcoma^33^, and diffuse large B-cell lymphoma (DLBCL)^34^. In a Phase 1 study of patients with desmoid tumours (NCT03459469), Tegavivint demonstrated favourable tolerability and promising therapeutic efficacy, achieving an overall response rate of 25% and a 9-month progression-free survival rate of 79%^35^. Recently, several phase 1/2 studies have initiated for patients with lymphomas and desmoid tumours (NCT04851119), leukaemia (NCT04874480), osteosarcoma (NCT07144254), diffuse large B-cell lymphoma (NCT05755087), lung non-small cell carcinoma (NCT04780568) and hepatocellular carcinoma (NCT05797805). Preclinical efficacy of Tegavivint has, as yet, not been described.

A range of preclinical models of hepatocellular carcinoma (HCC) are available, including *in vivo* models in several species, including mice. However, an important initial step in the validation of pathway efficacy in HCC is testing in human liver cancer cell lines. HepG2, HUH6, and HUH7 are three human-derived liver cancer cell lines that are frequently used, well characterised and exhibit differing levels of Wnt/β-catenin signalling activation. These cell lines are not derived from hepatitis B virus (HBV)-or hepatitis C virus (HCV)-infected tumours. In addition, HepG2 and Huh7 cells can be transplanted into immunodeficient mice to generate xenograft models for *in vivo* drug efficacy testing^36^.

Here, we describe a systematic analysis of the effect of Tegavivint effect on human liver cancer-derived cell lines. We also evaluated safety and anti-cancer efficacy of Tegavivint treatment in an orthotopic xenograft transplant mouse model using HepG2 cells.

## Results

### HepG2, HUH6 and HUH7 HCC cell lines exhibit a different level of activation of Wnt/β-catenin signalling

First, we compared activation of the Wnt/β-catenin pathway in HepG2, HUH6 and HUH7 HCC cell lines. The stability of β-catenin is regulated by a specific phosphodegron motif, DpSGXXpS (residues 32–37), within the β-catenin protein^9^. The priming kinase CK1α phosphorylates the Ser45 position, which triggers sequential phosphorylation by GSK3β at Thr41, Ser37, and Ser33 for degradation of β-catenin by β-Transducin repeat containing protein (β-TrCP). In human cancers, these specific residues constitute mutational hotspots^14^. We used the HepG2, HUH6 and HUH7 lines to represent a variety of both Wnt/β-catenin activated and non-activated cells: HepG2 has a large *CTNNB1* deletion including the mutational hotspots^37^; HUH6 has a point *CTNNB1* mutation creating a G34V substitution ^37^ in the phosphodegron motif; and HUH7 has no *CTNNB1* mutations^38^ (Fig. 1A). Following cell line authentication, we validated β-catenin pathway activation using immunofluorescence to detect nuclear β-catenin localisation (Fig. 1B), confirming strong activation in the HepG2 lines, weaker activation the HUH6 line and no activation in the HUH7 lines, consistent with previous reports^37–39^. Consistent with this, *AXIN2*, a robust marker for Wnt pathway activation^40^, *NOTUM*, a specific marker for Wnt pathway activation^41, 42^, and *LGR5*, less specific than *AXIN2* as also functioning as a tissue stem cell marker^43^, were expressed at higher levels in the HepG2 lines, compared to HUH6 and HUH7 lines^39^ (Fig. 1C). Expression of *AXIN2* and *LGR5* in the HUH6 lines indicated activation of the Wnt pathway in this cell line. *LGR5*, *AXIN2* and *NOTUM* were low in the HUH7 line consistent with inactive β-catenin pathway^39^, but this lines does express GS as reported previously^44^. Since *GLUL* expression can be regulated by mechanisms independent of Wnt signalling^45^, it may not serve as a reliable readout for pathway activity in HUH7 cells.

**Fig. 1.**
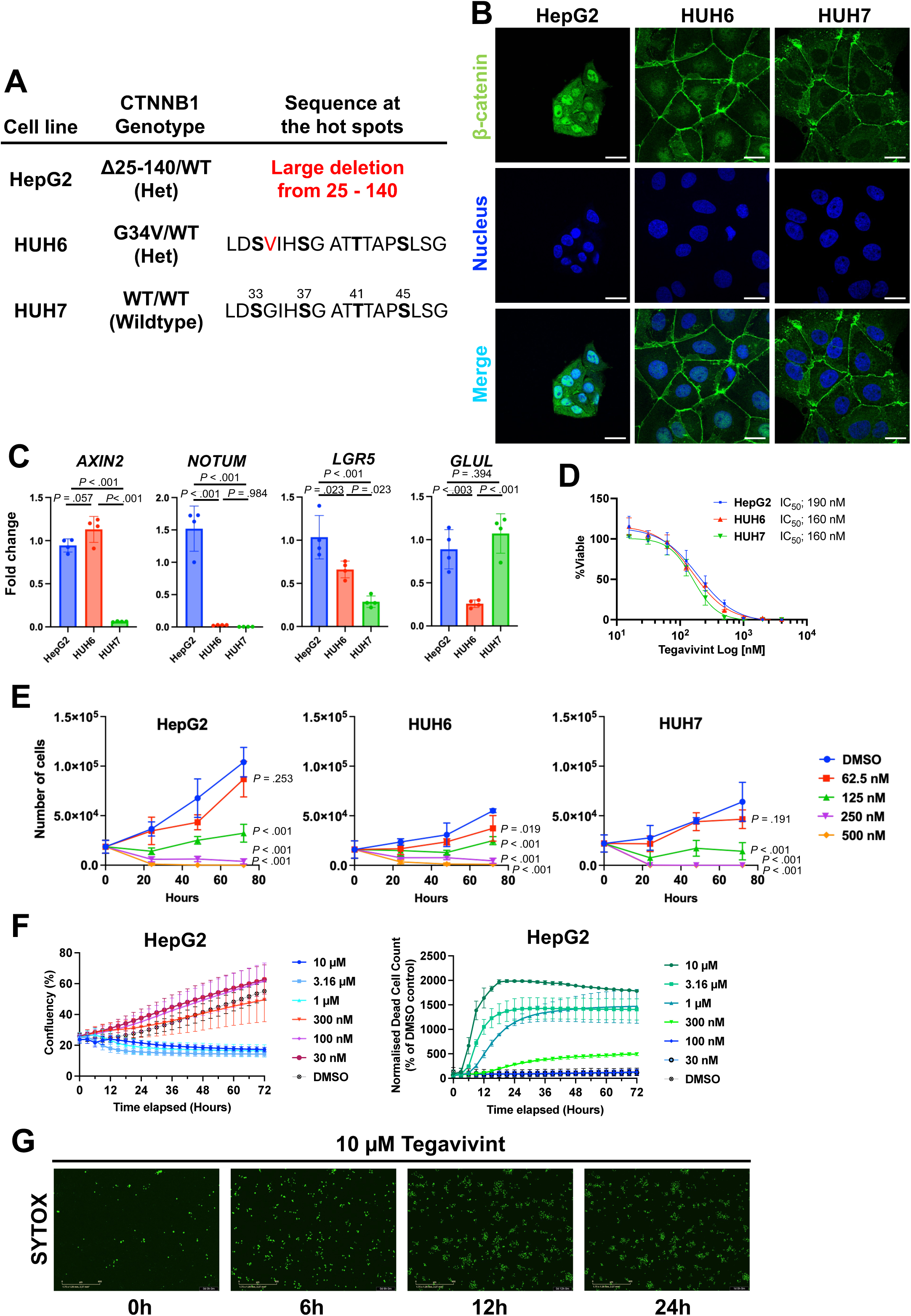
Tegavivint induces cell death in liver cancer cells. (**A**) β-catenin (*CTNNB1*) status of liver cancer cell lines. Phosphorylation sites, bold. Mutations, red. (**B**) Representative images of immunofluorescence for β-catenin (green) and nucleus (blue). Scale bars, 20 µm. Two repeated experiments with two technical replicates per experiment (**C**) Fold change in expression levels (relative to RPS18 housekeeping; normalised to HepG2). Each dot represents an experiment (n=4 repeated experiments). Bars are mean±SD. Data were analysed with one-way ANOVA with Tukey’s multiple comparisons test. (**D**) Cell viability was assessed following 48 hours of treatment with Tegavivint. Data were normalised to DMSO (0.025%). Each data point represents the mean±SEM of n=3 repeated experiments with three technical replicates per experiment. (**E**) Cells were treated with Tegavivint at indicated concentrations or DMSO (0.0031%) as a control. Total viable cell counts were determined at indicated hours post-treatment using manual counting. Data points represent the mean±SD of n=3 repeated experiments with no technical replicate per experiment. Data were analysed with one-way ANOVA followed by Dunnett’s multiple comparisons test on the 72-hour time point to compare each treatment group against the DMSO control. (**F**) Growth of HepG2 cells was monitored using the Incucyte Live-Cell Analysis System. Data are percentage (%) of confluence (Left) and cell death as SYTOX Green^+^ cells (Right). Mean±SD; n=3 repeated experiments. (**G**) Representative images of SYTOX Green. Scale bars, 400 µm.

### Tegavivint, a novel class of Wnt/**β**-catenin pathway inhibitor, shows high cytotoxic potency to human liver cancer cell lines *in vitro*

To test the effect of Tegavivint on human liver cancer cell lines, we performed dose-response studies. Each cell line was then treated with up to 4 µM Tegavivint *in vitro* and response was assessed by ATP bioluminescence assay to quantify the live cell number after 48 hours. Each cell line was consistently killed by Tegavivint with IC_50_ values of 190, 160 and 160 nM in HepG2, HUH6 and HUH7 lines, respectively (Fig. 1D). There was some evidence of reduced potency in the β-catenin pathway active HCC lines at higher doses compared with the HUH7 line at this 48-hour timepoint. We, therefore, went on to test over a 3-day time course with a variety of doses of Tegavivint in all 3 cell lines (Fig. 1E). Negative controls, treated with only DMSO, grew as expected over time with HepG2 cells undergoing the largest expansion. When treated with increasing doses of Tegavivint, the number of cells in all three lines was markedly reduced at 500 nM, consistent with our 48-hour dose response finding. Interestingly, growth suppression was maintained and more pronounced in the HUH7 line than both the HUH6 and HepG2 β-catenin mutant lines. Notably, 250 nM Tegavivint suppressed but did not ablate the growth of these lines, whilst 125 nM inhibited at 24-48 hours but permitted ongoing growth to 72 hours. Therefore, Tegavivint potently kills all three liver cancer cell lines with no apparent relationship between cytotoxicity and the activation of the Wnt/β-catenin pathway.

We then proceeded to explore the kinetics of cell growth and death in the prototypic β-catenin mutant HepG2 line. Cells were monitored kinetically using SYTOX™ Green to visualize cell death via the Incucyte™ live-cell analysis system. Untreated HepG2 cells grew, but Tegavivint treatment caused a dose dependent growth suppression at doses above 300 nM. Examining cell death, doses above 100 nM produced cell death in the HepG2 line, with higher doses producing earlier cell death within the first 24 hours (Figs. 1F and G).

### Tegavivint treatment causes cell cycle arrest in HepG2 and HUH7 cells but not in HUH6 cells

Given the role of β-catenin in promoting cell proliferation in both hepatocytes and liver cancer cells, we explored the effects of Tegavivint upon cell cycle in the three liver cancer cell lines. We utilised a range of Tegavivint doses *in vitro*, from 62.5-500 nM; spanning the IC_50_ values and doses that effectively suppress survival in all three lines. Analysis of cell cycle distribution 24 hours post-treatment revealed that higher doses of Tegavivint consistently induced transition from G1 into a quiescent G0 state (determined as Ki-67 negative cells) (Figs. 2A and B). Similarly, at higher doses, both HepG2 and HUH7 cells exhibited a decreased proportion of cells in G1 phase, accompanied by a moderate accumulation in the G2 phase. Consistent with the observed G2 accumulation, both cell lines showed a significant reduction in the mitotic (M phase, determined as phospho-Histone H3 positive cells) fraction, suggesting a pre-mitotic arrest. (Fig. 2C) with the β-catenin mutant cell lines appearing most responsive at lower doses of Tegavivint below the IC_50_ (e.g. 62.5 nM).

**Fig. 2.**
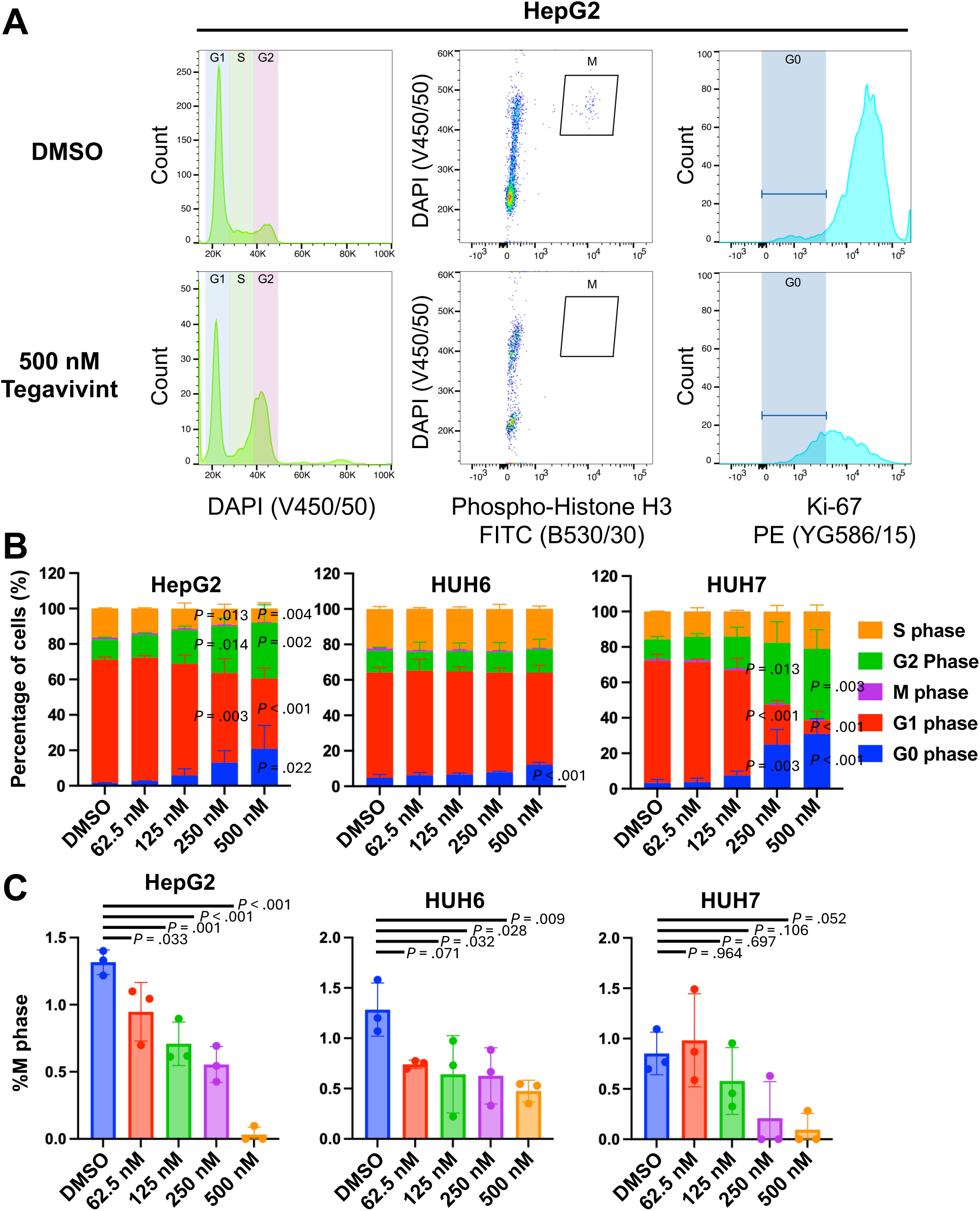
Tegavivint induces cell death in liver cancer cells. (**A**) Representative flow cytometry plots of HepG2 cell line treated with DMSO (0.0031%) or 500 nM Tegavivint-. Total DNA content was assessed by DAPI intensity (Left). Mitotic cells were identified by phospho-Histone H3^+^ and DAPI^High^ (Middle), and G0 cells were marked by Ki67^-^ cells (Right). (**B**, **C**) Percentage of HepG2, HUH6 or HUH7 cells in G0, G1, G2/M and S phases following treatment with DMSO (0.0031%) or indicated concentrations of Tegavivint for 24 hours. M-phase percentages are shown as bar graph in C. Data represent the mean±SD of n=3 repeated experiments. Data were analysed by one-way ANOVA followed by Dunnett’s multiple comparisons test, with each cell cycle phase compared to DMSO control.

### Tegavivint treatment induces apoptosis in HUH6 cells and non-apoptotic cell death in HepG2 and HUH7 cells

Next, given the evidence of cell death, previous reports of apoptotic cell death^29–32^ and apoptosis independent cell death^46^, we explored the cell death status of each cell line at 24 hours post therapy. A proteasome inhibitor MG-132 was used as the positive control for induced apoptosis^47^. Interestingly, HUH6 cells treated with MG-132 exhibited greater sensitivity to apoptosis, as evidenced by an increase in phosphatidylserine-positive live cells (early apoptosis) and cleaved caspase-3-positive live cells (Figs. 3A–E). HepG2 cells showed an intermediate sensitivity compared to HUH6 cells and HUH7 cells. Surprisingly, MG-132 treatment of HUH7 cells did not induce apoptosis, as the majority of dead cells were cleaved caspase-3-negative (Fig. 3D). Tegavivint treatment highly induced apoptosis in HUH6 cells, as evidenced by an increase in cleaved caspase-3-positive live cells (Fig. 3D), while the majority of cell death in HUH7 cells was cleaved caspase-3-negative. HepG2 cells also exhibited both cleaved caspase-3-positive and -negative cell death.

**Fig. 3.**
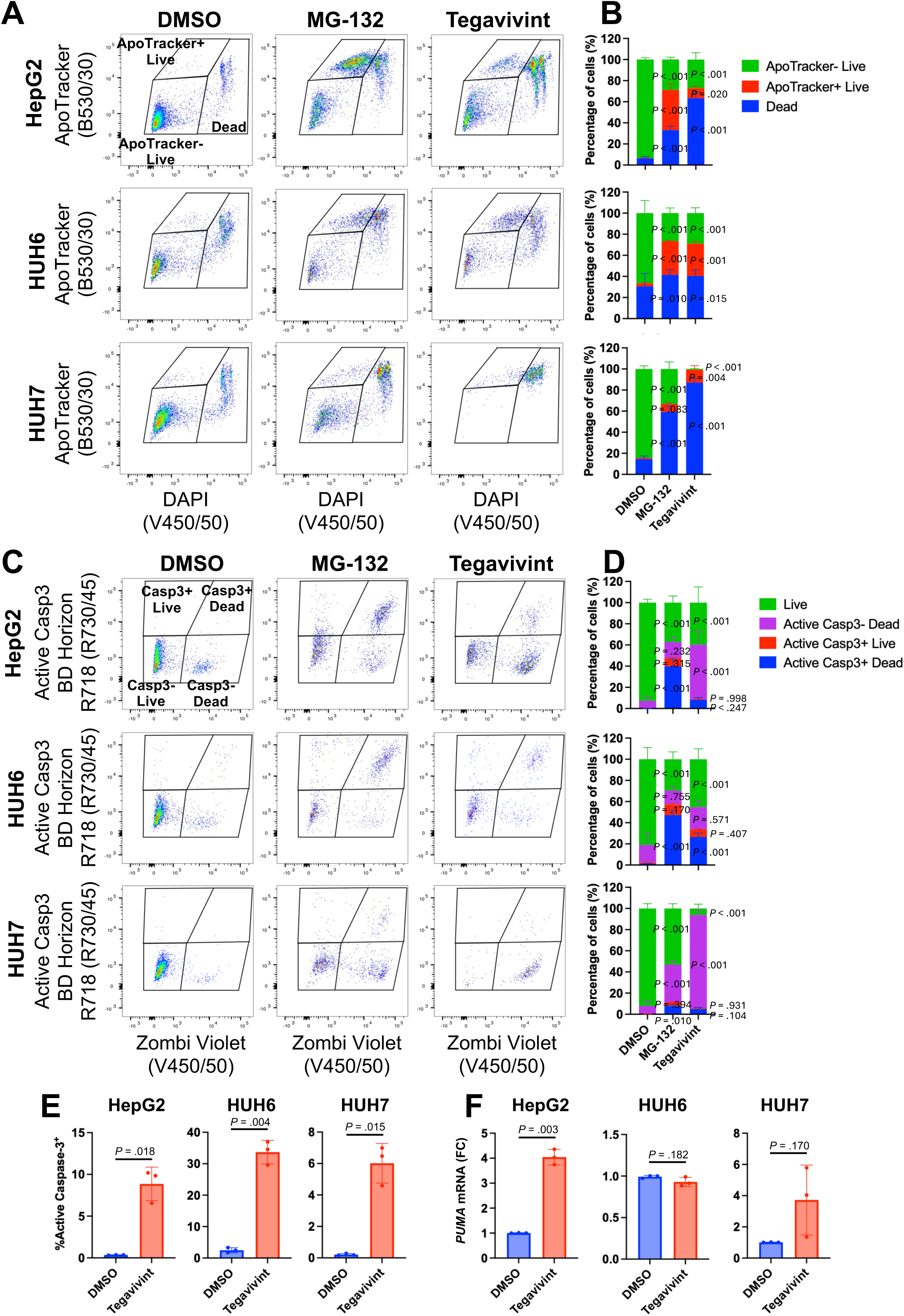
Apoptosis in liver cancer cells in response to Tegavivint. (**A**, **B**) Representative flow cytometry plots of indicated cell lines treated with DMSO (0.0031%), 50 µM MG-132, or 500 nM Tegavivint for 24 hours. Bar graphs representing the percentage of live cells (Apotracker^-^/DAPI^-^), ApoTracker^+^ live cells (early apoptotic cells, Apotracker^+^/DAPI^-^), and dead cells (dead cells, Apotracker^+^/DAPI^+^) in indicated cell lines following treatment for 24 hours. Data represent the mean±SD of n=3 repeated experiments. Data were analysed by one-way ANOVA followed by Dunnett’s multiple comparisons test, with each population compared to DMSO-treated control cells. **(C**, **D**) Representative flow cytometry plots of indicated cell lines treated with DMSO (0.0031%), 50 µM MG-132, or 500 nM Tegavivint for 24 hours. Bar graphs representing the percentage of live cells (Caspase-3^−^/Zombie^−^), active casp3^-^ dead (non-apoptotic dead cells, Caspase-3^−^/Zombie^+^), active casp3^+^ live cells (early apoptotic cells, Caspase-3^+^/Zombie^−^), and active casp3^+^ dead cells (late apoptotic cells, Caspase-3^+^/Zombie^+^). Data represent the mean±SD of n=3 repeated experiments. Data were analysed by one-way ANOVA followed by Dunnett’s multiple comparisons test, with each population compared to DMSO-treated control cells. (**E**) The total percentage of active caspase-3-positive cells (Caspase-3^+^/Zombie^−^ and Caspase-3^+^/Zombie^+^ populations) is presented in the bar graph. Unpaired t-test with Welch’s correction. (**F**) *PUMA* (pro-apoptotic) mRNA expression levels measured in the indicated cells treated with either DMSO or 500 nM Tegavivint for 16 hours by quantitative PCR and normalised to Ribosomal protein S18 (*RPS18*). E-F Bars are mean±SD of n=3 repeated experiments. Data were analysed by unpaired t-test with Welch’s correction.

Interestingly, these results together with cell cycle analysis revealed an inverse correlation between cell cycle regulation and apoptosis across the three lines (Cell cycle arrest, HUH7>HepG2>HUH6; Apoptosis, HUH6>HepG2>HUH7). HUH7 cells exhibited the most robust cell cycle arrest in response to Tegavivint, which corresponded with the lowest sensitivity to apoptosis. Conversely, HUH6 cells showed minimal effect on cell cycle but the highest rate of apoptotic induction. HepG2 showed an intermediate characteristic. These data suggest that HUH7 cells prioritise checkpoint-mediated survival and potential repair, whereas HUH6 cells bypass prolonged cell-cycle arrest in favour of immediate pro-apoptotic signalling. Therefore, the differences of phenotypes per cell line by Tegavivint treatment derives from the cell line characteristics, and Tegavivint has the potential to induce apoptosis as total cleaved caspase-3-positive cells (live and dead) were significantly increased in all cell lines treated with Tegavivint (Fig. 3E). Upregulation of the pro-apoptotic pathway (*PUMA*) was pronounced in the HepG2 line (Fig. 3F). This may indicate that apoptotic signalling can be activated but has comparative resistance to execute apoptosis in HepG2 versus the HUH6 tumour line.

### Tegavivint treatment suppress Wnt target genes in Wnt-activated liver cancer cell lines HepG2 and HUH6 cells

Then we explored on target activity of Tegavivint by measuring Wnt/β-catenin responsive genes in cells after 16 hours after Tegavivint therapy. We used 500 nM Tegavivint to explore this as this dose lies between the separation in growth seen in the HepG2 cells when measured longitudinally (Fig. 1F). Exploring a panel of Wnt/β-catenin target genes (Figs. 4A-E), we found no suppression of the expression of Wnt targets measured in the HUH7 cells. Interestingly, this included the expression of *GLUL* encoding glutamine synthetase (GS), implying that the expression of GS is not dependent upon Wnt/β-catenin signalling in HUH7 cells. *LGR5*, *AXIN2* and *NOTUM* were highly expressed by HepG2 (Fig. 1C), and Tegavivint suppresses expression of *LGR5* and *NOTUM* (Figs. 4A and C), along with an additional Wnt/β-catenin target gene, *Cyclin D1* (*CCND1*) (Fig. 4E). HUH6 cells have an intermediate level of *LGR5* and high expression of *AXIN2* (Fig. 1C), and Tegavivint treatment mildly suppresses *AXIN2* expression (Fig. 4B). As HUH6 cells were highly susceptible to apoptosis, we believe it is more challenging to test an ideal time-window to observe downregulation of Wnt/β-catenin target genes. There was no detectable suppression of *GLUL* in any of the cell lines. Overall, these results demonstrate that Tegavivint suppresses the Wnt/β-catenin pathway in Wnt/β-catenin activated liver cancer cell lines. Tegavivint may also utilise a Wnt/β-catenin-independent killing mechanism in HUH7 HCC cells.

**Fig. 4.**
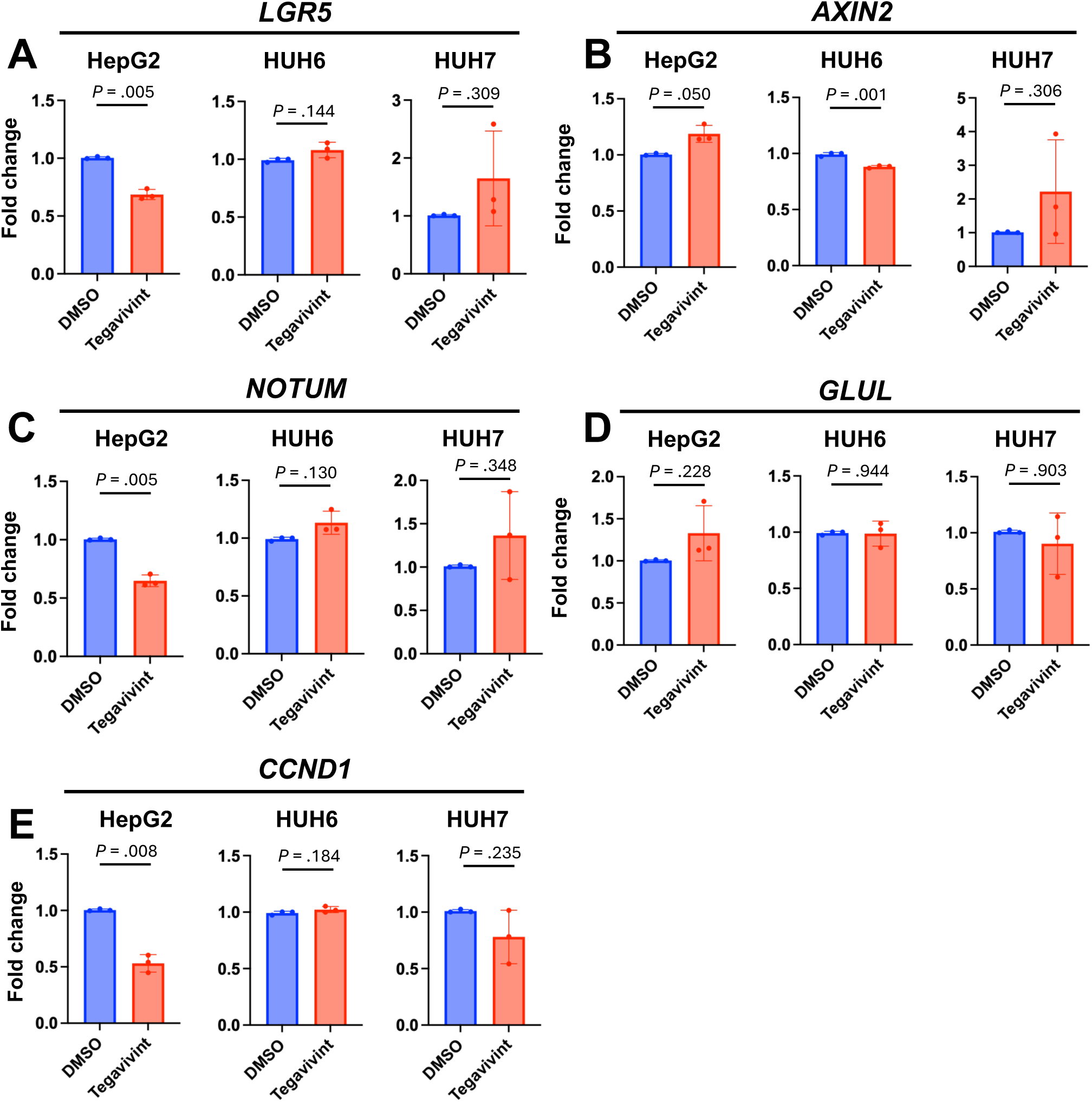
Assessment of the Wnt/β-catenin pathway suppression *in vitro*. (**A**-**F**) HepG2, HuH6 and HuH7 cells were treated with either DMSO or 500 nM Tegavivint for 16 hours. Relative mRNA expression levels of the indicated genes was measured by quantitative PCR and normalised to the expression of Ribosomal protein S18 (*RPS18*). Bars are mean±SD of n=3 repeated experiments. Statistical significance was determined by an unpaired t-test with Welch’s correction.

### Tegavivint suppresses tumour growth without major adverse effects in mice

Given the effects of Tegavivint upon the human liver cancer cell lines, we expanded our analysis to explore its potential for *in vivo* use of Tegavivint by using a humanised murine orthotopic liver tumour model. To model systemic administration of Tegavivint for the therapy of liver tumours, we set about to develop an orthotopic xenotransplantation model^48^. For this, we surgically implanted HepG2 cells into the left lobes of the liver in NOD-SCID immunodeficient adult male mice. Following recovery, mice were monitored clinically until day 14, when Tegavivint was administered systemically via intravenous injection, twice weekly. Mice were then harvested 3 weeks later whilst on treatment and compared to vehicle treated controls (Fig. 5A).

**Fig. 5.**
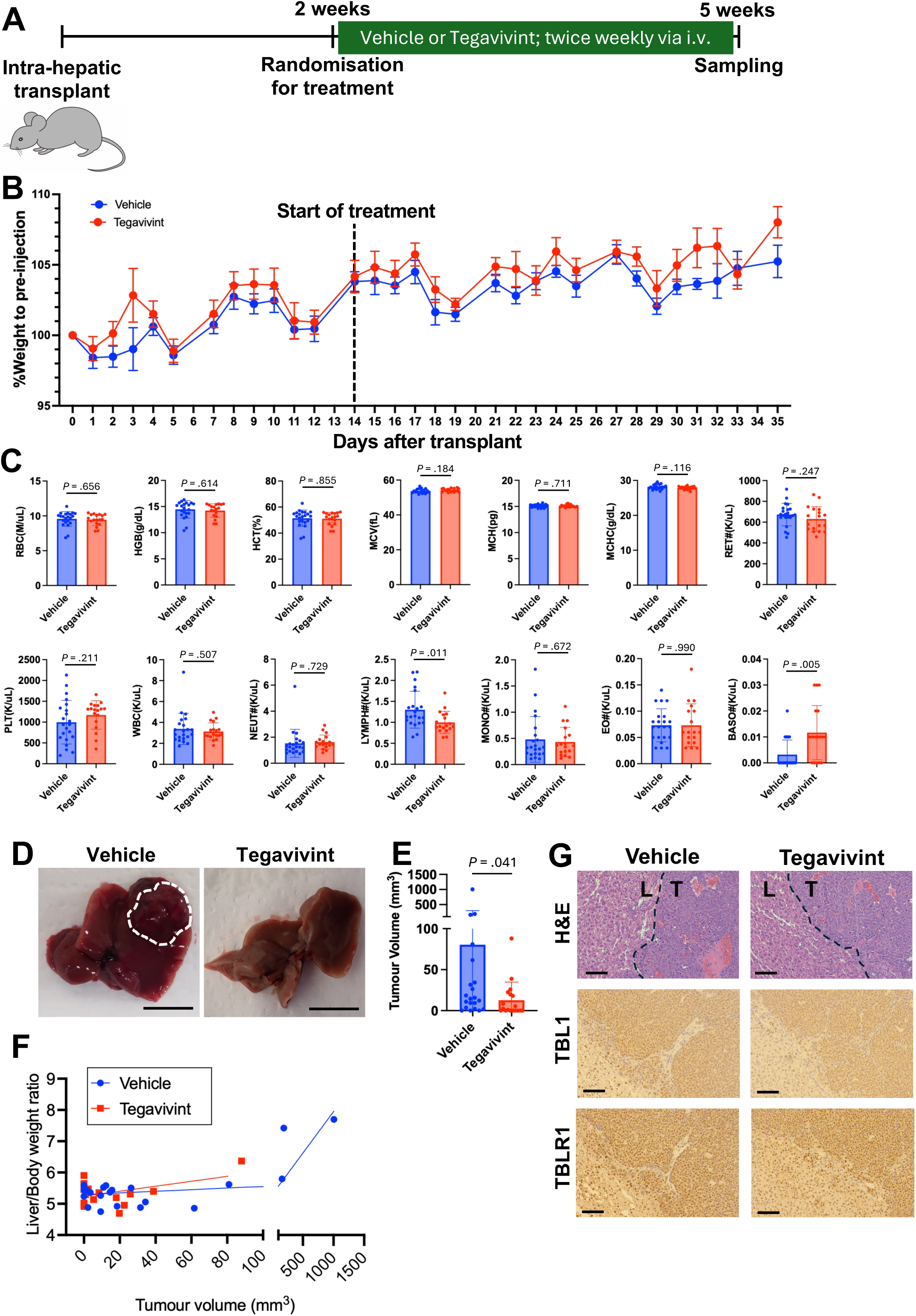
Humanised orthotopic model of Tegavivint therapy *in vivo*. (**A**) Schematic of the orthotopic transplant experiment. HepG2 cells were injected into the left liver lobe. Mice were randomly assigned to receive either vehicle (5% dextrose) or Tegavivint intravenously twice weekly from 2-weeks post transplantation. (**B**) Body weight changes relative to the day of injection. Data points represent the mean±SD of mice from three repeated experiments: Experiment 1 (Vehicle, n=5; Tegavivint, n=5), Experiment 2 (Vehicle, n=9; Tegavivint, n=9), and Experiment 3 (Vehicle, n=8; Tegavivint, n=8), totalling 44 mice). The weight was not significantly changed between the Vehicle or Tegavivint treated mice at any time point; The weight on Days 8–10 and 14–35 was significantly (P < 0.05) altered compared to Day 0. (**C**) Haematological parameters of mice following vehicle (n=22) or Tegavivint (n=18) treatment were measured five weeks post-transplantation using an IDEXX automated haematology analyser. Bars are the mean±SD of mice from three independent experiments (n=22 vehicle or n=18 Tegavivint). Data were analysed by unpaired t-test with Welch’s correction. (**D**) Representative images of the liver at sampling. The white dotted line delineates the liver tumour. Scale bar, 1 cm. (**E**) Quantification of total tumour burden (mm^3^) at 5 weeks post-intrahepatic injection (vehicle, n=21; Tegavivint, n=18). Data were analysed by a Mann-Whitney test. Bars are the mean±SD. (**F**) Correlation between liver weight-to-body weight ratio (%) and total tumour burden (mm^3^); lines represent linear regression (R^2^= 0.604 and 0.201 for vehicle and Tegavivint respectively; slope≠0 <0.0001 for vehicle). (**G**) Representative images of H&E staining and immunohistochemistry for TBL1 and TBL1R (brown) in transplanted HepG2 (human) tumours (T) and surrounding healthy (mouse) liver (L) treated with vehicle (n=4) or Tegavivint (n=3). Scale bar, 50 µm.

All mice remained well without adverse clinical signs following orthotopic transplantation and therapy in both treatment arms. There was no evidence of weight loss in either arm in the transplanted mice (Fig. 5B) nor complications from surgery in the mice treated with Tegavivint. Blood sampling was performed at the endpoint. There were no significant differences in circulating full blood counts between the study arms, with the exception of mild reductions in lymphocytes and increases in basophils. Notably, lymphocyte counts were expectedly low given the immunodeficient nature of these animals (Fig. 5C).

Tumours were assessed macroscopically at the predefined timepoint, and liver weight relative to body weight was measured to assess tumour volume. Tumour volume was variable in the model but was reduced in mice that received Tegavivint (Figs. 5D and E). Equally, there was a trend towards reduction in the liver weight in the Tegavivint treated arm, which acted as an independent surrogate for the size of the liver tumour as expected (Fig. 5F). Importantly, these tumours expressed both TBL1 and TBLR1, which are required for the activity of Tegavivint (Fig. 5G).

### Tegavivint suppresses Wnt target genes and proliferation to induce apoptosis in tumours in mice

Finally, to assess on target activity of Tegavivint, we examined canonical β-catenin transcriptional targets in both the orthotopic liver tumours and the background murine liver in each treatment group. As tumour material was sparce in the active treatment group and to ensure that signals were being explored in the tumours themselves, we relied principally on spatially resolved protein/transcript detection within the tumoural tissues. There was an apparent reduction of total cellular β-catenin following Tegavivint therapy in the orthotopic liver tumours, both in the cytoplasm and the nucleus (Fig. 6A). *LGR5* expression was also reduced in the orthotopic HepG2 tumours of mice treated systemically with Tegavivint compared to vehicle control (Fig. 6B and C). The transplanted tumours continued to express low levels of *GLUL* (encoding glutamine synthetase), which was reduced at transcript level in tumours of mice treated with Tegavivint compared to those treated with vehicle (Fig. 6D). Cell proliferation, visualized by human Ki-67 staining, was suppressed in tumours treated with Tegavivint (Figs. 6E and F), consistent with *in vitro* analysis. The number of TUNEL-positive cells was significantly increased in tumours treated with Tegavivint (Figs. 6G and H), Again consistent with *in vitro* data demonstrating induction of apoptosis.

**Fig. 6.**
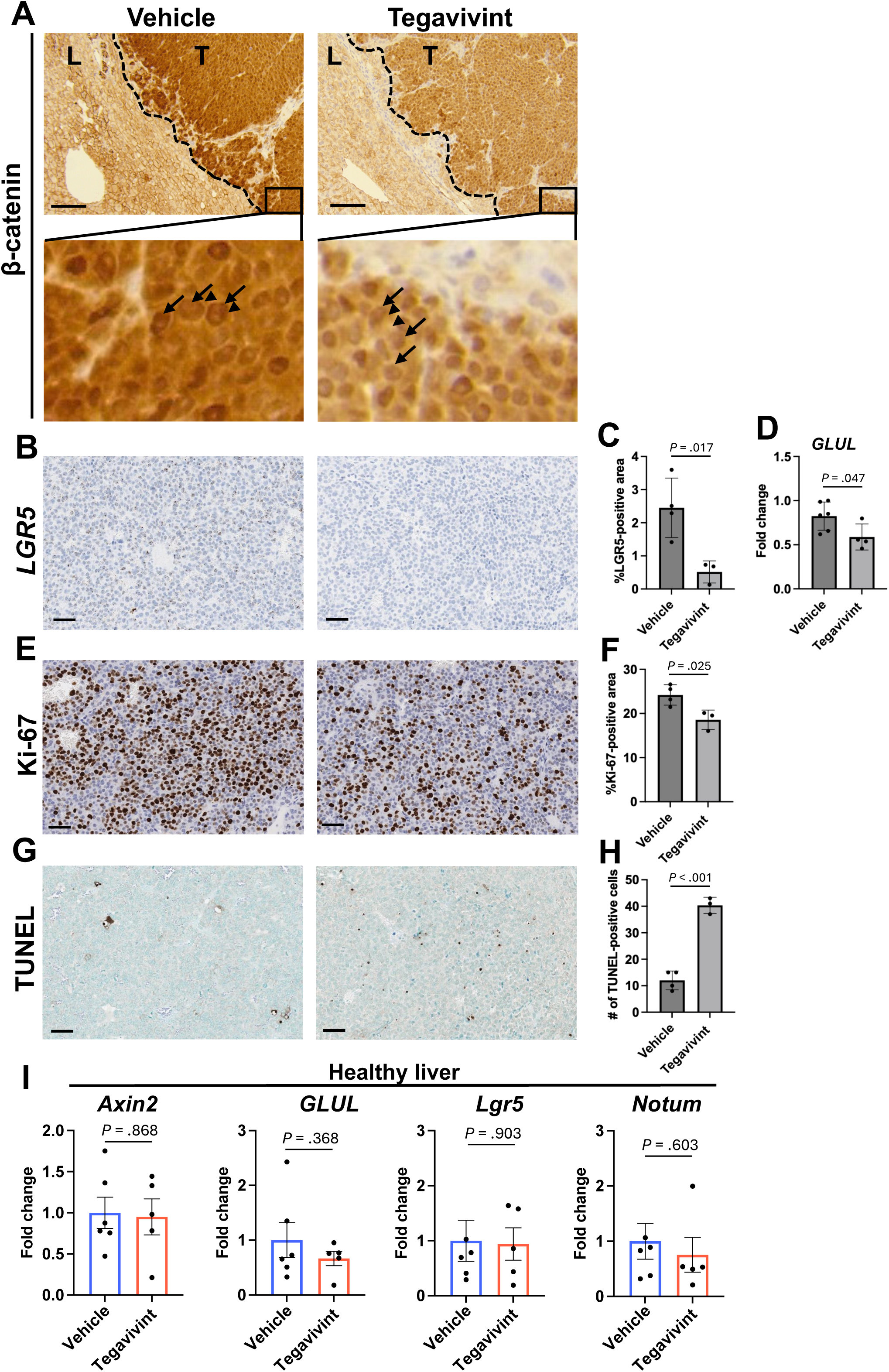
On target Wnt/β-catenin pathway suppression *in vivo*. Representative images of immunohistochemistry for β-catenin (Nuclei and cytoplasm are indicated by arrows and arrowheads, respectively.) (**A**) and human Ki-67 (**E**), *in situ* hybridisation for human *LGR5* mRNA (**C**), and TUNEL staining (**G**) in HepG2-forming tumours (T) and surrounding healthy mouse liver (L) treated with vehicle (n=4) or Tegavivint (n=3). Scale bar, 50 µm. (**B**) Relative *GLUL* mRNA expression levels were measured by quantitative PCR and normalised to Ribosomal protein L13A (*RPL13A*) in HepG2-forming tumours treated with vehicle (n=6) or Tegavivint (n=4). (D,F,H) Percentage of positively stained area per field of view (FOV): *LGR5* mRNA (D), human Ki-67 (F) and TUNEL (H). Bars are mean±SD; vehicle (n=4) or Tegavivint (n=3). Data were analysed with an unpaired t-test with Welch’s correction. (**I**) Indicated murine genes in the normal livers of mice treated with vehicle (n=6) or Tegavivint (n=5) and normalised to murine *Rps18*. Bars are mean±SD. Data were analysed with an unpaired t-test with Welch’s correction.

Examining the expression of Wnt/β-catenin target genes using mouse specific primers in the non-tumoural liver tissue, we did not find evidence of Wnt/β-catenin pathway’s suppression in the background liver (Fig. 6I). This suggest that the inhibition of the Wnt/β-catenin pathway by Tegavivint is comparatively specific for human tumours with minimal effects on normal murine tissue.

In summary, our *in vivo* findings demonstrate that Tegavivint effectively suppresses tumour growth in Wnt/β-catenin-driven models without eliciting detectable systemic toxicity or significant weight loss. These data establish Tegavivint as a potent and well-tolerated inhibitor of liver tumorigenesis, supporting its potential as a targeted therapeutic strategy for this molecular subtype (Wnt-activated subtype) of HCC.

## Discussion

In this study, we evaluated the effects of Tegavivint, a novel Wnt/β-catenin pathway inhibitor, on liver cancer both *in vitro* and *in vivo*. This is the first such study in liver cancer adding to prior descriptions of Tegavivint efficacy in preclinical mouse models of haematological malignancies^29,31,34^ and osteosarcoma^49^. Notably, Tegavivint inhibited the growth of all liver cancer cell lines tested, with an IC_50_ in the low hundred-nanomolar range, demonstrating its potent inhibitory activity against Wnt-activated liver cancer models. The tumour killing activity of Tegavivint (48-hour IC_50_ 190 nM in HepG2) was more potent than that of established Wnt/β-catenin antagonists such as XAV939 (7-day IC_50_ ∼80,000 nM in HepG2)^50^, PRI-724 (24-hour IC_50_ ∼20,000 nM in HepG2)^51^, and CGP049090 (72-hour IC_50_ 600 nM in HepG2)^52^, reinforcing its potential as a high-affinity therapeutic agent for Wnt-driven malignancies.

A recent study evaluated efficacy of β-catenin siRNA encapsulated in lipid nanoparticles (LNPs) using a preclinical mouse HCC model using forced expression of human mutant β-catenin^25^. The study demonstrated that high doses resulted in mortality, while moderate doses effectively suppressed tumour growth. In the present study, Tegavivint did not inhibit Wnt/β-catenin signalling in the healthy mouse liver, indicating its potential to avoid the on-target adverse effects on normal cells seen with β-catenin siRNA. Consistently, it has been reported that Tegavivint exhibits low cytotoxicity toward normal B cells^34^. Although the amino acid sequence homology of TBL1 and TBLR1 is high between humans and mice (human/mice correlation of 96% and 99% for TBL1 and TBL1R respectively - Protein BLAST search), species-specific differences in efficacy remain possible, as a less conserved region (aa 177–202 in NP_005638.1) preceding the WD40 domain may alter binding affinity to Tegavivint due to conformational differences. Furthermore, β-catenin siRNA therapy significantly inhibited tumours even in HCC models lacking CTNNB1 mutations^25^, suggesting that the Wnt/β-catenin pathway plays a broader role in HCC development. This may offer therapeutic potential of Tegavivint and other Wnt pathway inhibitors to a wider spectrum of HCC patients aside from CTNNB1 mutations alone. Inhibition of the β-catenin pathway using an siRNA approach activated interferon signalling, thereby inducing responsiveness to immune checkpoint inhibitors (ICIs)^25^ as implied from other preclinical studies^18^. In this present study, a xenograft NOD-SCID mouse model was utilized as an *in vivo* model to accommodate a human liver cancer cell line with CTNNB1 mutation. Consequently, the analysis of interactions with immune cells was limited; future studies using such immune-proficient mouse models driven by human β-catenin mutant are encouraged to explore potential of enhancing the efficacy of combination therapies with ICIs such as atezolizumab and bevacizumab. We elected not to persue organoid models in this study as organoid culture media typically contain Wnt agonists, which lead to constitutive activation of the Wnt/β-catenin pathway, regardless of the *CTNNB1* mutation status.

To elucidate the mechanism of action of Tegavivint, we investigated its effects on the cell cycle and apoptosis. First, positive control experiments using the proteasome inhibitor MG-132 revealed that two cell lines (HUH7 and HepG2) exhibited comparative resistance to apoptosis while the HUH6 line is more sensitive to apoptosis. In the apoptosis-sensitive HUH6 cell line, Tegavivint strongly induced apoptosis. In apoptosis-resistant HUH7 and HepG2 cells, Tegavivint increased the expression of *PUMA*, a marker of apoptosis; however, it did not strongly induce classical apoptosis characterized by cleaved caspase-3^53^. Instead, Tegavivint prominently induced cell cycle arrest in these cells. Subsequently, non-apoptotic pathways were likely activated, ultimately leading to cell death. One possible mechanism may involve lipid-dependent non-apoptotic cell death, as recently reported^46^, however, further studies are required to clarify how Tegavivint promotes cell cycle arrest and induces non-apoptotic cell death.

Tegavivint is known to exert its effects by inhibiting the interaction between β-catenin and TBL1/TBL1R^29^. Consistent with previous reports in AML^29^ and multiple myeloma^31^, This study showed that Tegavivint suppressed the expression of Wnt target genes (*LGR5*, *NOTUM*, and *Cyclin D1*) in HepG2 cells. In contrast, no changes in Wnt target gene expression were observed in HUH7 cells, which exhibit low activation of the Wnt/β-catenin pathway. This finding highlights a potential advantage of Tegavivint based on its unique mechanism of targeting TBL1/TBL1R. Other Wnt/β-catenin pathway inhibitors are known to suppress Wnt/β-catenin signalling even in normal cells, resulting in toxicity when administered *in vivo*. Because the Wnt pathway plays a critical role in normal intestinal regeneration and maintenance of bone density, preclinical and clinical trials showed that conventional Wnt inhibitors are associated with adverse effects such as intestinal and bone toxicity^54–57^. Since TBL1/TBL1R is more highly expressed in cancer cells than in normal cells^28^, Tegavivint may selectively target “Wnt-mutant” cancers that are addicted to Wnt signalling, thereby providing a more tumour-specific therapeutic strategy. Consistently, the phase I clinical trial of Tegavivint on desmoid tumours showed no severe adverse effects by Wnt inhibition using Tegavivint^35^.

Finally, we evaluated the safety and antitumor activity of Tegavivint *in vivo*. We chose the HepG2 cell line because availability of HCC cell lines with activated Wnt signalling is limited and it is a widely used liver cancer cell line, allowing comparison with previous studies of other Wnt inhibitors. Moreover, among the three cell lines tested, HepG2 exhibited clear activation of the Wnt pathway. To better recapitulate the unique hepatic microenvironment, we employed an orthotopic transplantation model in NOD-SCID mice. Consistent with our *in vitro* findings, decreased expression of Wnt target genes, reduced proliferation, and increased apoptosis were observed in tumours treated with Tegavivint. Tumour size was also markedly reduced. Because NOD-SCID mice exhibit profound immunodeficiency, the tumour regression observed in this study is unlikely to result from immune-mediated clearance and instead supports the direct induction of apoptosis in cancer cells by Tegavivint. Therefore, in immunocompetent contexts, Tegavivint may elicit even greater therapeutic efficacy, as the non-apoptotic cell death may trigger immunogenic cell death (ICD)^58, 59^, leading to activation of dendritic cells and T cells. Furthermore, in normal liver tissue, no reduction in Wnt target gene expression was observed. In addition, no significant changes in body weight or notable haematological toxicity were detected, suggesting that Tegavivint acts selectively on cancer cells. Although this study utilized a paediatric cancer–derived cell line (HepG2, derived from hepatoblastoma, Wnt mutations are also observed in 44% of adult hepatocellular carcinoma (HCC) cases^4^, suggesting that Tegavivint may be effective in a broad subset of HCC patients with strong Wnt pathway activation.

In conclusion, our study identifies Tegavivint as a potent inhibitor of the Wnt/β-catenin signalling pathway in liver cancer. By triggering apoptotic and non-apoptotic cell death pathways and demonstrating robust anti-tumour efficacy without systemic toxicity *in vivo*, Tegavivint addresses a critical unmet need for Wnt-driven malignancies. These findings provide a compelling preclinical rationale for the clinical development of Tegavivint as a targeted therapy for patients with Wnt-mutated hepatocellular carcinoma.

## Materials and methods

### Cell culture

HepG2, HUH6 and HUH7 HCC cell lines were generously provided by Dr. Saverio Tardito (Medical University of Vienna). All three are derived from male patients. HepG2 cells were cultured in Minimum Essential Medium (MEM) with 1 g/l glucose (21090022, Gibco) supplemented with 2 mM glutamine (25030-024, Gibco), 1% non-essential amino acids (11140035, Gibco), 1 mM pyruvate (S8636, Sigma-Aldrich), 10% foetal bovine serum (FBS) (A5256701, Gibco) and 1x Penicillin-Streptomycin (15140-122, Gibco). HUH6 and HUH7 cells were cultured Dubecco’s modified Eagle medium (DMEM) with 4.5 g/l glucose and sodium pyruvate (21969-035, Thermo Fisher Scientific), supplemented with 10% FBS, 2 mM glutamine, 1x Penicillin-Streptomycin.

Cells were maintained in T175 flasks and incubated at 37°C and 5% CO_2_. Cells were passaged using phosphate-buffered saline PBS containing 0.25% trypsin (15090-046, Gibco) and 0.5mM EDTA twice a week. Cells were seeded at their desired density in new flasks or plates (*in vitro*). Mycoplasma testing was routinely performed and cultures always tested negative for Mycoplasma contamination. HepG2, HUH6 and HUH7 cells were confirmed as authentic cell lines using the Promega GenePrint 10 Kit (B9510, Promega).

### Immunofluorescence staining

HepG2, HUH6, and HUH7 cells were seeded onto glass coverslips in 12-well plates at a density of 2 × 10 cells per well. After 24 h, cells were washed three times with ice-cold PBS and fixed with 4% paraformaldehyde in PBS (pH 7.4) for 15 min at room temperature. Coverslips were washed twice with PBS and permeabilised with 0.1% Triton X-100 in PBS for 3 min. Following three additional washes with PBS (5 min), cells were blocked for 30 min in 1% bovine serum albumin (BSA) in PBS containing Tween-20 (PBS-T). Cells were incubated with anti-β-catenin primary antibody (1:100; BD Biosciences, 610154) diluted in blocking solution for 1 h at room temperature in a humidified chamber. After three washes with PBS (5 min), cells were incubated for 1 h at room temperature with Alexa Fluor 488-conjugated goat anti-rabbit IgG (H+L) secondary antibody (1:100; A-11008, Molecular Probes, Thermo Fisher Scientific) diluted in blocking solution. Coverslips were then washed twice with PBS and once with Milli-Q water, air-dried, and mounted onto glass slides using ProLong™ Diamond Antifade Mountant with DAPI (P36962, Invitrogen). Slides were stored at 4°C in the dark. Images were acquired using a Zeiss LSM 710 confocal microscope equipped with a Plan-Apochromat 63×/1.40 oil immersion DIC M27 objective. Representative images were processed using ImageJ software (v1.54g).

### Drug treatment

For *in vitro* experiments, Tegavivint (supplied by Iterion) and MG-132 (M8699, Sigma-Aldrich) were dissolved in dimethyl sulfoxide (DMSO) (10213810, Fisher Scientific) to prepare a 16 mM and 50 µM stock solution, respectively, stored at -20°C and diluted in cell culture medium to the desired working concentrations.

### RNA extraction, cDNA synthesis and quantitative PCR

For RNA was extraction extracted from cell lines, HepG2, HUH6, and HUH7 cells were seeded at a density of 200,000 cells per well in 6-well plates. 24 hours after plating, cells were treated with either DMSO or 500 nanomolar (nM) Tegavivint for 16 hours. Cells were washed with PBS once, frozen by dry ice and stored in aat -80oC freezer until lysis. 600 µL Buffer RLT was added directly into a well, scraped using a cell scraper, and homogenized homogenised using a QIAshredder (79654, Qiagen). For RNA was also extraction extracted from healthy tissues, a piece of liver and liver tumours and liver tissues upon dissection that were dissected and were frozen by dry ice and before stored in aage at -80oC freezer until lysis. 5 x 5 x 5 mm piece of frozen liver or tumour tissue was cut out using scissors and transferred lace into a soft tissue homogenizing homogenising CK14 2 mL tube (P000912-LYSK0-A, Precellys) containing 600 µL Buffer RLT from the RNeasy kit (74106, Qiagen), and kept on ice until homogenization homogenisation by a Precellys Tissue Homogenizer (programme, soft tissue). After homogenizationhomogenisation, RNA was isolated using the RNeasy kit (74106, Qiagen). RNA concentration and purity was were determined using a NanoDrop spectrophotometer. cDNA was synthesized synthesised from 500 ng RNA using a Quantitect Reverse Transcription Kit (205311, Qiagen).

Quantitative PCR was performed using a QuantiTect SYBR Green PCR Master Mix (204145, Qiagen) on a QuantStudio™ 3 real-time PCR system (A28131, Thermo Fisher Scientific). PCR cycle settings were: denaturation at 94°C for 15 s, annealing at 55°C for 30 s and extension at 72°C for 30 s; repeated for 45 cycles. Melt curve analysis and data analysis were performed on a QuantStudio design and analysis software (v.1.5.2). Relative expression was calculated by the ΔC_t_ method using Ribosomal protein S18 as an endogenous control. The following primer sequences were used for each gene: Ribosomal protein S18 (*RPS18*) forward 5’-GATGGGCGGCGGAAAATAGC-3’, reverse 5’-CGCCCTCTTGGTGAGGTCAA-3’; Cyclin D1 (*CCND1*) forward 5’-GGCGGAGGAGAACAAACAGA-3’, reverse 5’-GGAGGGCGGATTGGAAATGA-3’; PUMA (*BBC3*) forward 5’-CCAAACGTGACCACTAGCCT-3’, reverse 5’-GATGAAGGTGAGGCAGGCAT-3’. The other primers were purchased from QIAGEN (Quantitect Primer): human LGR5 (QT00027720); human NOTUM (QT00018025); human AXIN2 (QT00037639); human GLUL (QT00085155); human RPL13A (QT00089915); murine Lgr5 (QT00123193); murine Notum (QT01749559); murine Axin2 (QT00126539); murine Glul (QT01062306); murine Rn18s (QT02448075).

### Cell viability assay

HepG2, HUH6, and HUH7 cells were seeded at a density of 10,000 cells per well in 96-well plates. 24 hours after plating, cells were treated with either DMSO or Tegavivint at 15.625, 31.25, 62.5, 125, 250, 500, 1000, 2000 or 4000 nM for 48 hours. Cell viability was assessed using the CellTiter-Glo® Luminescent Cell Viability Assay (G7572, Promega) following the manufacturer’s instructions. Luminescence was measured on a GloMax® Navigator microplate luminometer (GM2010, Promega). Results were normalised to the vehicle control. Curve fitting and IC_50_ values were determined using a nonlinear regression model (Sigmoidal, 4-parameter logistic regression).

### Cell growth curve assay

HepG2, HUH6, and HUH7 cells were seeded at a density of 20,000 cells per well in 24-well plates. Twenty-four hours after plating, cells were treated with either DMSO or Tegavivint in 500 µL of medium for 24 hours. At 24, 48, and 72 hours after treatment, the culture medium was removed, and each well was washed with 500 µL PBS. Cells were then detached by adding 100 µL PBS containing 0.25% trypsin and 0.5 mM EDTA and incubating for 7 minutes at 37°C. Cells were dissociated into single cells by gentle pipetting. 5 µL of cell suspension was mixed with 5 µL trypan Blue (15250-061, GIBCO), and counted on a haemocytometer (Brand).

### Real-time proliferation and death assays

Cell proliferation and cytotoxicity were monitored in real-time using the Incucyte® Live-Cell Analysis System (Sartorius). HepG2 cells were seeded at a density of 7,500 cells in 96-well plates and allowed to adhere overnight. Following treatment, the culture medium was supplemented with SYTOX™ Green (S7020, Thermo Fisher Scientific) at a final concentration of 60 nM to detect loss of membrane integrity. Plates were placed in the Incucyte® incubator, and images were captured every 3 hours for a total duration of 72 hours. Cell growth was quantified as a percentage of phase area confluence. Cell death was measured as the SYTOX-positive count normalised by the count of DMSO-treated wells. All data were analysed using the Incucyte® AI Cell Health Analysis software module.

### Flow cytometry

HepG2, HUH6, and HUH7 cells were seeded at a density of 20,000 cells per well in 24-well plates. Twenty-four hours after plating, cells were treated with either DMSO or Tegavivint in 500 µL of medium for 24 hours. The culture medium was collected, and each well was washed with 500 µL PBS, which was also collected. Cells were then detached by adding 100 µL PBS containing 0.25% trypsin and 0.5 mM EDTA and incubating for 7 minutes at 37°C. Cells were dissociated into single cells by gentle pipetting and collected. The collected medium, PBS wash, and trypsinised cells were combined and centrifuged at 400 × g for 5 minutes. The supernatant was discarded, and cells were resuspended in 0.5% BSA (A7906, Sigma-Aldrich) in PBS for cell cycle or apoptosis analysis.

For cell cycle analysis, cells were resuspended in 100 µL of 0.5% BSA in PBS, followed by the addition of 100 µL IC Fixation Buffer (00-8222-49, Thermo Fisher Scientific). After thorough mixing, cells were incubated for 10 minutes at room temperature. Cells were centrifuged at 400 × g for 5 minutes and the supernatant was discarded. 100 µL ice-cold 100% methanol was added and mixed well, followed by incubation for at least 30 min at 4°C. 100 µL 0.5% BSA in PBS was added, and centrifuged at 400 × g for 5 minutes and the supernatant was discarded. Cells were resuspended in 100 µL 0.5% BSA in PBS containing anti-human/mouse Ki-67 Antibody conjugated with PE (130-120-557, Miltenyi Biotec) at 1:500 dilution, anti-human/mouse Histone H3 pS28 Antibody conjugated with FITC (130-105-699, Miltenyi Biotec) at 1:100 dilution and 5 ug/ml DAPI (422801, BD), and incubated for 30 minutes at room temperature under dark. 100 µL 0.5% BSA in PBS was added and centrifuged at 400 × g for 5 minutes and the supernatant was discarded. Cells were resuspended in 0.5% BSA in PBS before acquisition on flow cytometer.

For phosphatidylserine detection, cells were resuspended in 100 ul 0.5% BSA in PBS containing 400 nM stock Apotracker Green (427402, BioLegend) and incubated at room temperature for 10 minutes. 100 µL 0.5% BSA in PBS containing 5 ug/ml DAPI was added and centrifuged at 400 × g for 5 minutes and the supernatant was discarded. Cells were resuspended in 0.5% BSA in PBS before acquisition on a BD LSRFortessa flow cytometer.

For active Caspase 3 detection, cells were resuspended in 200 µL cold PBS, centrifuged at 400 × g for 5 minutes, and the supernatant was discarded. Cells were resuspended in 100 µL Zombie Violet (423113, BioLegend) at 1:100 in cold PBS, incubated for 20 minutes at 4 °C. 100 µL 0.5% BSA in PBS was added, and centrifuged at 400 × g for 5 minutes and the supernatant was discarded. Cells were resuspended in 100 µL IC fixation buffer and incubate 20 minutes at 4 °C. 100 µL permeabilization buffer (00-8333-56, Thermo Fisher Scientific) was added and centrifuged at 400 × g for 5 minutes and the supernatant was discarded. Cells were resuspended in 100 µL permeabilisation buffer containing anti-Active Caspase-3 conjugated with BD Horizon™ R718 (570311, BD) at 1:100 dilution and incubated for 30 minutes at 4 °C. 100 µL permeabilisation buffer was added and centrifuged at 400 × g for 5 minutes and the supernatant was discarded. Cells were resuspended in 200 ul 0.5% BSA in PBS and centrifuged at 400 × g for 5 minutes and the supernatant was discarded. Cells were resuspended in 0.5% BSA in PBS before acquisition on a BD LSRFortessa flow cytometer with DIVA software (BD Biosciences). Data were analysed using FlowJo Software version 10.10.0.

### Mouse experiment/transplant

All mouse experiments were performed in accordance with a UK Home Office project licence (PP0604995; protocol number 8), in accordance with Home Office guidelines, and were subject to review by the animal welfare and ethical review board of the University of Glasgow. To minimise pain, suffering and distress to the animals, single-use needles and non-adverse handling techniques were used throughout. The mice used in this study were male NOD-SCID mice (commercially bought from Charles River Laboratories, UK or Inotiv, Netherlands). Mice were housed in a specific high barrier environment and kept under standard conditions with a 12h day/night cycle with unrestricted access to food and water ad libitum. Environmental enrichments, in the form of bedding, plastic tunnels and chew sticks, were added to all cages. Plastic tunnels were removed from cages for the first 7-10 days after orthotopic transplant to avoid catching/tearing of the skin clips.

For cell preparation for injection, HepG2 cells were detached using PBS containing 0.25% trypsin and 0.5 mM EDTA, and medium was added, and centrifuged at 180 × g (1,000 rpm) for 5 minutes. Cells were washed with cold PBS, and then 100,000 cells were resuspended in 10 µL cold Corning Matrigel and kept on ice until injection. Age- and batch-matched commercially bought male mice were given a minimum 1-week acclimatisation period before orthotopic intrahepatic injection between 8 and 13 weeks of age. All mice were administered 150 μL subcutaneous supportive fluids (PBS) in the flank and analgesia prior to surgery [5 mg/kg Carprofen (Rimadyl) in the drinking water from 24 h before surgery and for a further 48 h after surgery], and 0.03 mg/ml subcutaneous administration of Buprenorphine (Vetergesic) in the scruff. Mice were anaesthetised using sevoflurane (with O_2_) in a sterile surgical field. A median laparotomy was performed followed by expression of the left lateral liver lobe. 100,000 HepG2 cells in 10 μL of Corning Matrigel were injected into the left lobe of the liver using a Hamilton syringe. Sterile surgical gauze was placed over the injection site to prevent blood loss and leakage of cells. The liver was gently placed back inside the abdomen, and the muscle layer was closed with a continuous vicryl suture (Ethicon). A local anaesthetic [2.5 mg/ml Bupivacane (Marcaine)] was topically administered to the sutured wound prior to skin clipping. Hydrogel H_2_O and diet gel packs (Tecniplast) were added to the cage during recovery. Mice were placed in a warmed recovery cage to assist recovery for up to 4 hours. Surgical clips were removed between 7 and 10 days post-transplantation. All mice were health checked daily and were housed in the same location. Tegavivint was formulated in nanoparticle form and suspended in 5% dextrose and stored at room temperature. Tegavivint or 5% dextrose was administered to mice at 50 mg/kg (60 µL) intravenously twice weekly before randomisation at 2 weeks post-transplantation (https://www.graphpad.com/quickcalcs/randomize1/).

### Dissection and scoring of tumour burden

Mice were humanely euthanised at the 5-week time point by CO inhalation using a rising concentration method, followed by cervical dislocation. Blood was collected by cardiac puncture and placed in EDTA-coated collection tubes (Sarstedt). Following dissection, whole livers were weighed then further dissected into individual liver lobes, and visible macroscopic liver tumours were scored using digital callipers in either two (<1 cm) or three dimensions (≥1 cm) for calculating tumour volume (length/2×height/2×breath/2×4/3π). The liver/tumour were fixed in 10% neutral buffered formalin (in PBS) for 24 h, followed by replacement with 70% ethanol.

### Haematology analysis

Blood was collected by cardiac puncture at the 5-week time point, and approximately 100 µL was transferred into EDTA-coated tubes and analysed using an IDEXX ProCyte Dx haematology analyser (IDEXX Laboratories, Westbrook, ME, USA) to measure haematological parameters.

### Histology

Dissected mouse tissue was fixed overnight in 10% neutral buffered formalin prior to transfer to 70% ethanol. Tissue was then processed with a standard overnight histology tissue-processing cycle before being orientated and embedded in histology wax. All Immunohistochemistry (IHC), in-situ hybridisation (ISH), Haematoxylin and Eosin (H&E) staining and TUNEL staining were performed on 4µm formalin fixed paraffin embedded sections (FFPE) that had previously been ovened at 60 C for 2 hours.

The following antibodies were stained on a Leica Bond Rx autostainer, Β-catenin (610154, BD Biosciences), Cleaved Caspase 3 (CC3) (9661, Cell Signaling Technology), TBL1X (66955-1-Ig, Proteintech) and TBLR1 (ab190976, Abcam). All FFPE sections underwent on-board dewaxing (AR9222, Leica) and epitope retrieval using appropriate retrieval solutions. Sections for TBL1X staining were retrieved using ER1 solution (AR9961, Leica) for 40 minutes at 100°C. Sections for B-Catenin, CC3 and TBLR1 staining underwent epitope retrieval using ER2 solution (AR9640, Leica) for 30 minutes (B-Catenin) and 20 minutes (CC3,TBLR1) at 100°C. Sections were rinsed with Leica wash buffer (AR9590, Leica) before peroxidase block was performed using an Intense R kit (DS9263, Leica) for 5 minutes. Sections were rinsed with wash buffer before sections for Β-catenin and TBL1X staining had mouse Ig blocking solution applied (MKB-2213, Vector Labs) for 20 minutes. Sections were rinsed with wash buffer before primary antibody application at an optimal dilution (B-Catenin, 1/250; CC3, 1/500; TBL1X, 1/3000; TBLR1, 1/100). The sections were rinsed with wash buffer before sections being stained for Β-catenin and TBL1X had mouse envision (K4001, Agilent) applied and sections for CC3 and TBLR1 staining had rabbit envision (K4003, Agilent) applied for 30 minutes. The sections were rinsed with wash buffer, visualised using DAB and counterstained with haematoxylin in the Intense R kit.

FFPE sections for Glutamine Synthetase (GS) (HPA-007316, Sigma-Aldrich) and Ki-67 (M7240, Agilent) staining were loaded into an Agilent pre-treatment module to be dewaxed and undergo heat induced epitope retrieval (HIER) using appropriate target retrieval solution (TRS) where sections were heated to 97 C for 20 minutes. Sections for Ki-67 staining underwent epitope retrieval using High pH TRS (K8004, Agilent) and sections for GS were retrieved using Low pH TRS (K8005, Agilent). After HIER the sections were rinsed in flex wash buffer (K8007, Agilent) prior to being loaded onto the Agilent autostainer. The sections underwent peroxidase blocking (S2023, Agilent) for 5 minutes and rinsed with flex buffer. The primary antibody application was at an optimised dilution (GS, 1/800; Ki-67, 1/100). Sections were washed with flex buffer before application of appropriate secondary antibody. Sections for Ki-67 staining had mouse envision and sections for GS had rabbit envision applied for 30 minutes. Sections were rinsed with flex wash buffer before applying Liquid DAB (K3468, Agilent) for 10 minutes. Sections were washed in water and counterstained with haematoxylin z (RBA-4201-00A, CellPath).

ISH detection for Hs-*LGR5* (311028), Mm-*Ppib* (313918) and *dap*β (312038) (Bio-Techne) mRNA was performed using RNAScope 2.5 LSx (Brown) detection kit (322700; Bio-Techne) strictly following the manufacturer’s instructions.

A TUNEL assay kit HRP-DAB (ab206836, Abcam) was used to stain apoptotic nuclei in the tissue sections strictly following manufacturer’s instructions.

To complete the IHC, ISH and TUNEL staining sections were rinsed in tap water, dehydrated through a series of graded alcohols and placed in xylene. The stained sections were coverslipped in xylene using DPX mountant (SEA-1300-00A, CellPath).

### Image analysis

Stained slide sections were scanned at 20× magnification (resulting in a resolution of 0.503 μm/pixel) using a Leica Aperio AT2 slide scanner. Scanned images were analysed blindly using HALO Image analysis software (V3.6.4134.396; Indica Labs). Representative areas of staining within the tumour were selected, and the staining-positive area within a defined region was quantified for LGR5 and Ki-67 using the Area Quantification v2.4.2 algorithm of the HALO software. For TUNEL, the number of TUNEL-positive puncta was manually counted on the HALO software within a defined region.

### Statistical analysis

Data are presented as mean±SEM or mean±SD, as indicated in the figure legends. Statistical analyses were performed using GraphPad Prism (version 10.6.1; GraphPad Software, San Diego, CA, USA). Differences between groups were assessed using appropriate statistical tests depending on the data type, as specified in the figure legends. A p value < 0.05 was considered statistically significant. For in vitro experiments, the sample size was determined using the ClinCalc online power analysis tool (https://clincalc.com/stats/samplesize.aspx). With α=0.05 and 80% power, n=3 independent experiments per group were required to detect a 50% treatment effect, assuming a 20% standard deviation (SD) in the control group. For in vivo experiments, power analysis (α=0.05, 80% power) indicated that n=16 animals per group were required to detect a 50% effect size, assuming a 50% SD in controls. No data were excluded with the exception of the orthotopic transplant experiments. For these, three independent experiments were performed: Experiment 1 (Vehicle, n=5; Tegavivint, n=5), Experiment 2 (Vehicle, n=9; Tegavivint, n=9), and Experiment 3 (Vehicle, n=8; Tegavivint, n=8), totalling 44 mice. In the second experiment, two mice were excluded: one due to surgical complications and weight loss (euthanised three days after the first Tegavivint dose), and another found dead on the day of the final dose (cause of death undetermined by post-mortem). Additionally, three samples were excluded because tumour areas could not be determined (one from Experiment 2 and two from Experiment 3). Finally, three samples (Vehicle, n=1; Tegavivint, n=2) were excluded from the analysis for Figures 5E and 5F due to technical measurement errors.

## Abbreviations

AML: acute myeloid leukaemia
BASO: Basophils
DLBCL: diffuse large B-cell lymphoma
EO: Eosinophils
FBS: foetal bovine serum
FFPE: formalin-fixed paraffin-embedded
FITC: fluorescein isothiocyanate
GS: glutamine synthetase
H&E: haematoxylin and eosin
HCT: Haematocrit
HGB: Haemoglobin
HIER: heat-induced epitope retrieval
ICD: immunogenic cell death
IHC: immunohistochemistry
ISH: in-situ hybridization
LYMPH: Lymphocytes
MAPK: mitogen-activated protein kinase
MCV: Mean Corpuscular Volume
MCH: Mean Corpuscular Haemoglobin
MCHC: Mean Corpuscular Haemoglobin Concentration
MEM: Minimum Essential Medium
MONO: Monocytes
NEUT: Neutrophils
NOD-SCID: non-obese diabetic/severe combined immunodeficiency
PLT: Platelets
RBC: Red Blood Cell
RET: Reticulocyte
TRS: target retrieval solution
TUNEL: terminal deoxynucleotidyl transferase dUTP nick end labelling
WBC: White Blood Cell.

## Acknowledgements

We thank the CRUK Scotland Institute’s Core Facilities (RRID:SCR_027384), in particular, Biological Services, Histology, Flow Cytometry Service, Molecular Technology (RRID:SCR_027368) and the Beatson Advanced Imaging Resource (RRID:SCR_023875) for their help. We thank Catherine Winchester (CRUK Scotland Institute) for advice on research integrity and manuscript reviewing.

## Authors’ contributions

Conceptualization: T.S., C.C., T.D., A.D., S.H., T.G.B.; Data curation: T.S., C.C., T.D., S.M., M.E.K., T.G.B; Formal analysis: T.S., C.C., M.E.K., T.G.B.; Funding acquisition: T.S., T.D., T.G.B.; Investigation: T.S., C.C., T.D., S.M., K.L.Y.S., A.G., M.Q., F.C., E.P., A.D., S.H., M.E.K., C.N., V.H.V, T.G.B.; Methodology: T.S., C.C., T.D., S.M., A.D., S.H., M.E.K., C.N., V.H.V, T.G.B.; Project administration: T.S., T.D., A.D., S.H., T.G.B.; Resources: A.D., S.H., T.G.B.; Supervision: T.S., T.G.B.; Validation: T.S., C.C. T.G.B.; Visualization: T.S., C.C., M.E.K., T.G.B.; Writing– original draft: T.S., C.N., T.G.B.; Writing – review & editing: T.S., T.G.B. All authors have approved the final version.

